# Dynamics of mechanically coupled hair-cell bundles of the inner ear

**DOI:** 10.1101/2020.08.20.259507

**Authors:** Y. Roongthumskul, J. Faber, D. Bozovic

## Abstract

The high sensitivity and effective frequency discrimination of sound detection performed by the auditory system rely on the dynamics of a system of hair cells. In the inner ear, these acoustic receptors are primarily attached to an overlying structure which provides mechanical coupling between the hair bundles. While the dynamics of individual hair bundles have been extensively investigated, the influence of mechanical coupling on the motility of the system of bundles remains underdetermined. We developed a technique of mechanically coupling two active hair bundles, enabling us to probe the dynamics of the coupled system experimentally. We demonstrated that the coupling could enhance the coherence of hair bundles’ spontaneous oscillation as well as their phase-locked response to sinusoidal stimuli, at the calcium concentration in the surrounding fluid near the physiological level. The empirical data were consistent with numerical results from a model of two coupled nonisochronous oscillators, each displaying a supercritical Hopf bifurcation. The model revealed that weak coupling can poise the system of unstable oscillators closer to the bifurcation by a shift in the critical point. In addition, the dynamics of strongly coupled oscillators far from criticality suggested that individual hair bundles may be regarded as nonisochronous oscillators. An optimal degree of nonisochronicity was required for the observed tuning behavior in the coherence of autonomous motion of the coupled system.

**STATEMENT OF SIGNIFICANCE:** Hair cells of the inner ear transduce acoustic energy into electrical signals via a deflection of hair bundles. Unlike a passive mechanical antenna, a free-standing hair bundle behaves as an active oscillator that can sustain autonomous oscillations, as well as amplify a low-level stimulus. Hair bundles under physiological conditions are elastically coupled to each other via an extracellular matrix. Therefore, the dynamics of coupled nonlinear oscillators underlie the performance of the peripheral auditory system. Despite extensive theoretical investigations, there are limited experimental evidence that support the significance of coupling on hair bundle motility. We develop a technique to mechanically couple hair bundles and demonstrate the benefits of coupling on hair bundle spontaneous motility.

## INTRODUCTION

Signal transduction by the inner ear is performed by a system of hair cells. Owing to the nonlinearity and internal active processes of hair cells, the inner ear displays exquisite sensitivity and high frequency resolution [1, 2]. Under constant mechanical forces, the apical protrusion of a hair cell, called the hair bundle, displays a nonlinear force-displacement relationship due to the gating of the mechano-sensitive transduction channels [3, 4]. The hair bundle under a sinusoidal driving force produces mechanical feedback which facilitates the amplification of low-level signals near a characteristic frequency [4 – 7]. In non-mammalian species, active bundle motility likely constitutes the primary source of amplification. In the mammalian cochlea, a process termed electromotility, which refers to the hair cell’s somatic length change upon variations in the membrane’s potential [8], also influences the frequency selectivity as well as the amplitude threshold of hearing [9, 10].

Hair bundle motility is well described by a nonlinear oscillator near a bifurcation: at the critical point, the stability of the system switches from a stationary fixed point to a self-sustained limit cycle. Under appropriate *in vitro* environments, undriven hair bundles robustly exhibit spontaneous oscillations [11], a behavior that corresponds to a limit cycle of a dynamical system. Under different types of chemical and mechanical manipulations on hair bundles of *in vitro* preparations, these autonomous oscillations can be enhanced, attenuated, or suppressed via a variety of bifurcations, including Hopf and saddle node on an invariant circle [12 – 14]. Near a critical point, individual hair bundles have been demonstrated to display enhanced sensitivity and frequency selectivity to sinusoidal stimuli [15, 16].

A dynamical system near a bifurcation also provides a generic phenomenological description for the inner ear [17, 18]. However, how the innate motility of an individual hair bundle contributes to *in vivo* phenomena of the inner ear remains unclear. In the inner ear of most species, hair bundles are tightly anchored to an overlying extracellular matrix, such as the tectorial membrane in the mammalian cochlea, or the otolithic membrane in the amphibian sacculus. This overlying structure provides strong coupling between neighboring bundles. Upon an increase in the bundles’ separation, the coupling strength becomes progressively weaker. The performance of the inner ear epithelium thus relies on the dynamics of hair cells under various degrees of mechanical coupling. Understanding the role of coupling would hence establish the connection between the dynamics of a hair bundle and the response of the full auditory system.

The dynamics of coupled nonlinear oscillators have been extensively investigated due to their relevance in various fields of study [19, 20]. For the auditory system, several theoretical investigations suggest benefits of coupling on signal detection performed by groups of active hair bundles. Theoretical predictions reveal that a system of oscillating hair bundles with a large frequency dispersion may become quiescent under a sufficiently strong coupling, a phenomenon termed amplitude death [21, 22]. This regime shows a greatly enhanced signal-to-noise ratio in response to a driving force, making it an attractive alternative mechanism that the auditory system may utilize to achieve sensitivity in the presence of noise. Further, coupling of unstable oscillators with a small difference in their characteristic frequencies reduces the effective noise level, resulting in more coherent autonomous oscillations, as well as the enhanced quality factor of their response to a sinusoidal driving force [23 – 26]. Although this has been experimentally demonstrated in a system of a small number of hair bundles [27, 28], the roles of coupling on hair bundle motility over a broader range of control parameters are underdetermined.

We investigate the dynamics of coupled hair bundles, both experimentally and theoretically. First, we study a mathematical model of two mutually coupled nonisochronous oscillators, each displaying a supercritical Hopf bifurcation. The theoretical predictions in the strong-coupling limit are used to describe the activity of two mechanically coupled hair bundles from the bullfrog sacculus. The bundles are brought across the critical point by increasing the level of calcium concentration of the artificial endolymph solution. Near the physiological calcium level, we observed two types of behavior of spontaneous dynamics as well as two types of response functions to sinusoidal forces of various frequencies. These distinct behaviors correspond to different degrees of nonisochronicity in the model of coupled oscillators. Finally, using a microscopic model of hair bundle mechanics, we illustrate that a calcium-sensitive elastic element within the bundle could potentially contribute to the nonisochronicity.

## MATERIALS AND METHODS

### Mathematical model of two mutually coupled nonlinear oscillators

The dynamics of each hair bundle was described by the normal form of a nonlinear system near a supercritical Hopf bifurcation (Results, Eqs. 1 – 2). The bundles’ positions were represented by the real parts of the complex variables *z*_1_(*t*) and *z*_2_(*t*). The imaginary parts served as hidden variables, not accessible in the experimental recordings. Both oscillators had identical control parameters *μ* and complex coefficients of nonlinearity, whose imaginary part *β* determined the degree of nonisochronicity. In the absence of coupling, a single oscillator resided at a stable fixed point when *μ* < 0, and displayed a limit cycle oscillation when *μ* > 0. The limit-cycle frequency Ω depended on the oscillation amplitude *R* and was determined by Ω = ω_0_ − *βR*^2^. The oscillation frequency of an isochronous oscillator, *β* = 0, was always equal to its characteristic frequency *ω*_0_.

The coupling term was modeled to facilitate comparisons to the physiological data. In our experiments, sets of two hair bundles were connected by attachment to the same glass fiber (Fig. 3A). The coupling stiffness between bundles due to an elastic force, estimated by the bending stiffness of a thin rod of a length equal to the bundle separation, was ∼1 N/m and significantly dominated the dissipative force (Supporting material, I). The coupling term was thus modeled as an elastic element whose stiffness was real and denoted by *K*. Its extension was determined by the difference in the oscillators’ displacements. We note that, in our model, direct coupling was limited to the real component and did not involve the imaginary part of the dynamical variables. In addition, the bundles-probe tip system was also coupled to the base of the fiber via a stiffness of ∼0.1 – 0.2 mN/m. This elastic element, however, was neglected in the model as it was significantly more compliant than the coupling between bundles.

**Figure 1.**
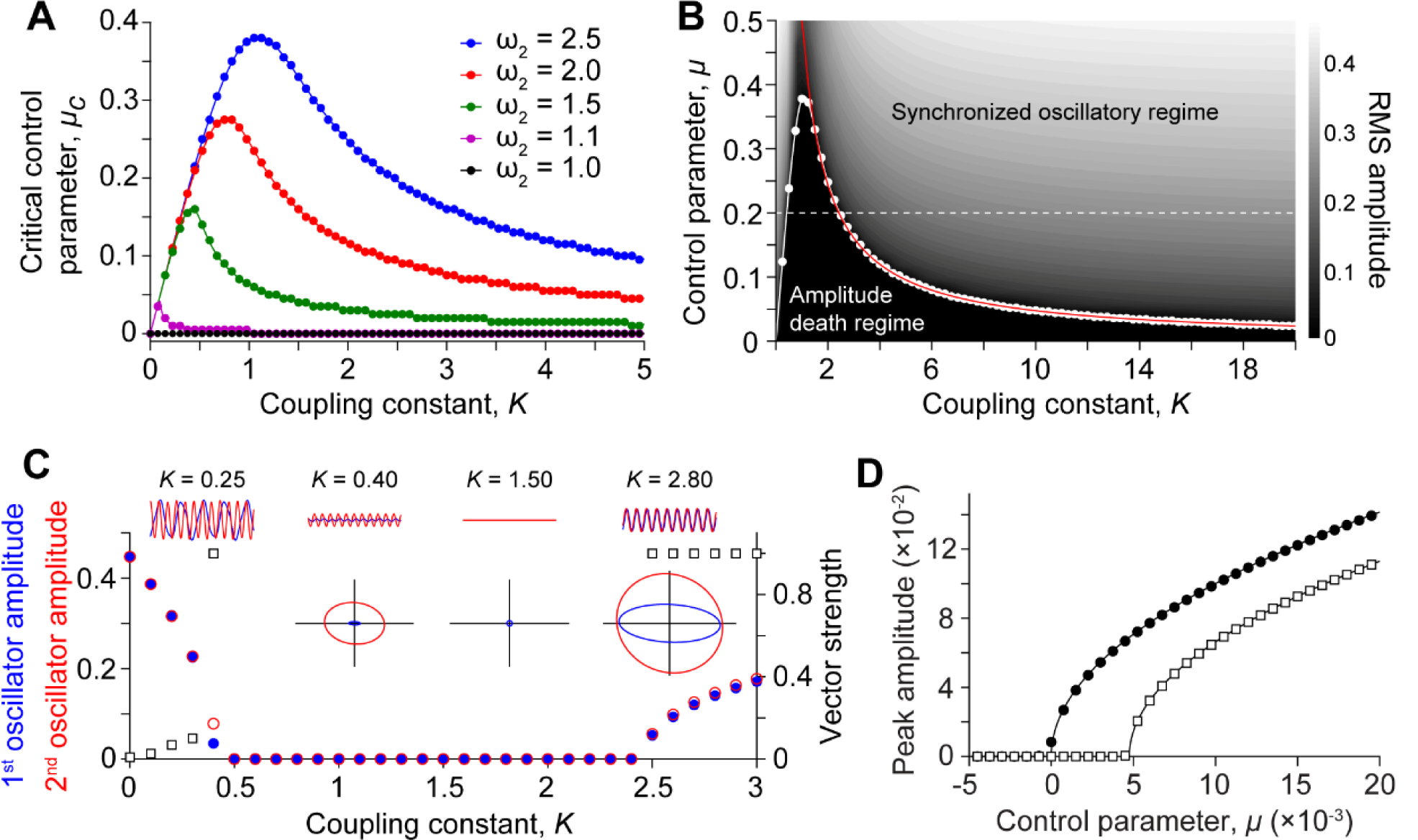
(*Model*) The critical control parameter (*μ*_*c*_) of coupled oscillators. In all panels, *ω*_0,1_ is fixed at 1. A) *μ*_*c*_ is estimated from the rate of amplitude change of the transient solution obtained from numerical simulations. The estimated *μ*_*c*_ remains at zero for identical oscillators: *ω*_0,1_ = *ω*_0,2_ = 1 (black dots) and progressively increases upon growth of the difference between the characteristic frequencies: *ω*_0,2_ = 1.1 (magenta), *ω*_0,2_ = 1.5 (green), *ω*_0,2_ = 2.0 (red), and *ω*_0,2_ = 2.5 (blue). *μ*_*c*_ calculated from a linear stability analysis of the model are consistent with the estimated values (solid lines). B) A state diagram of coupled oscillators with *ω*_0,2_ = 2.5 incorporates the line of supercritical Hopf bifurcation whose *μ*_*c*_ is estimated from numerical simulations (white dots), and calculated from Eq. 3 (white solid line). The heat map illustrates the RMS amplitude of the 1^st^ oscillator with brighter shades representing higher values. The oscillators are arrested when *μ* < *μ*_*c*_ (‘Amplitude death’). The vector strength of the oscillators’ phase difference reach unity over a regime with sufficiently large coupling constant or control parameter (‘Synchronized oscillation’). The onset of the regime is indicated by a red solid line interpolated from the smallest *K* that results in the unity vector strength obtained from numerical simulations. C) Same parameters as in B) with *μ* fixed at 0.2 (white dashed line in B)). The amplitudes of the 1^st^ (blue dots) and 2^nd^ (red open dots) oscillation vanish when *K*∼ 0.5 – 2.4. The vector strength of the phase difference (squares) indicates the absence of 1-to-1 phase locking for weak coupling and the onset of synchronization for *K* > 2.4. The time traces of both oscillations at different coupling constants illustrate the suppression of oscillatory activity (upper insets, blue lines: 1^st^ oscillator, red lines: 2^nd^ oscillator). The phase portraits at the corresponding coupling constant reveal a stable fixed point at the origin of the complex plane, an indication of an amplitude death regime (lower insets). D) At *K* = 100, the peak amplitude of the synchronized oscillation increases with the control parameter. *μ*_*c*_ is zero for a small frequency difference (*ω*_0,2_ = 1.1, black dots) and is positive for a large frequency difference (*ω*_0,2_ = 2.5, squares). Solid lines illustrate results from the fit: *μ*^1/2^ for *ω*_0,2_ = 1.1, and 0.915(*μ* − 4.75 × 10^−3^)^1/2^ for *ω*_0,2_ = 2.5.

**Figure 2.**
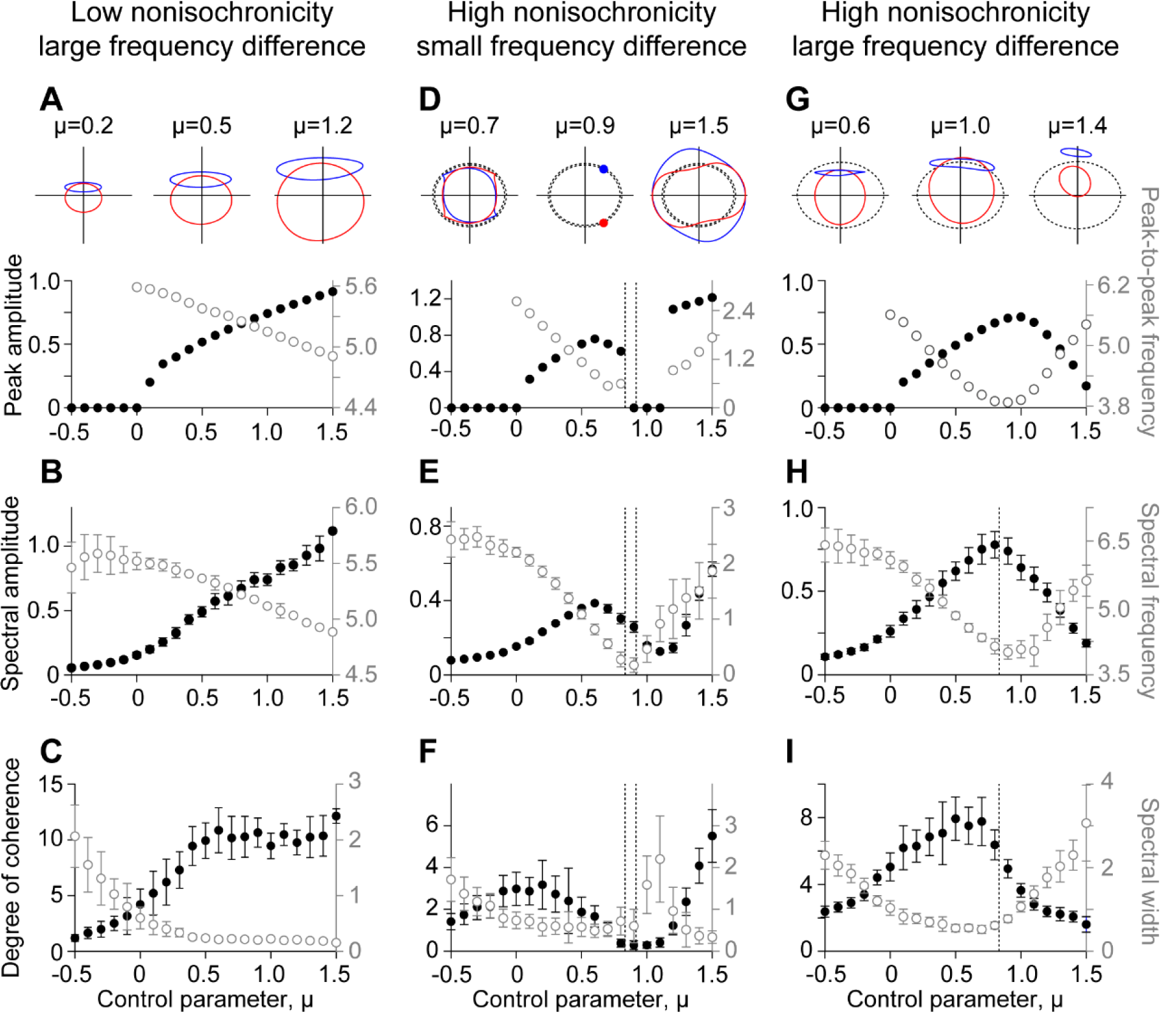
(*Model*) Dynamics of coupled oscillators far from criticality. The parameters are *ω*_0,1_ = 2.5, *K* = 100. In panels in which noise is included, the noise variance is *σ*^2^ = 4, and error bars indicate standard deviation of data obtained from 10 simulations. In A)-C), *β* = 0.5, and *ω*_0,2_ = 7.5. A) In the absence of noise, weakly nonisochronous oscillators display a monotonic increase in the peak amplitude of the oscillation (black dots), and a decline in the peak-to-peak frequency (gray open dots) upon growth of the control parameter in the oscillatory regime. (inset) The phase portraits at the corresponding control parameter reveal the distinct shapes of the limit cycles (blue lines: 1^st^ oscillator, red lines: 2^nd^ oscillator). B) When Gaussian white noise is included, the spectral amplitude (black dots) and spectral frequency (gray open dots) exhibit qualitatively identical behaviors as in the deterministic limit. C) The degree of coherence of the autonomous motion is enhanced (black dots) as the width of the spectral peak decreases (gray open dots) when the system enters the oscillatory regime. In D)-F), *β* = 3, and *ω*_0,2_ = 2.75. The vertical dashed lines indicate the control parameters at the angular nullclines of the 1^st^ and 2^nd^ oscillators (*μ*_0,1_ = 0.83 and *μ*_0,2_ = 0.92). D) Far from criticality, strongly nonisochronous oscillators with a small frequency difference are arrested in the absence of noise. (inset) The phase portraits illustrate the stable fixed points in the vicinity of the angular nullclines (dashed lines). E) With random fluctuations, the system exhibits tuned behavior. I) The coherence becomes minimal over a range of the control parameter coinciding with diminishing amplitude. In conjunction with this, the spectral peak reaches a maximum. In G)-I), *β* = 3, and *ω*_0,2_ = 7.5. The vertical dashed line indicates the control parameter at the angular nullcline of the 1^st^ oscillator (*μ*_0,1_ = 0.83). G) In the deterministic limit, strongly nonisochronous oscillators with a large frequency difference show a peak in the spectral amplitude and a minimum in the frequency near the angular nullcline. (inset) The limit cycles separate in association with the reduction of the oscillation amplitude, as the control parameter exceeds the value at the angular nullcline. H) Qualitatively similar behaviors are observed upon addition of noise. I) The degree of coherence exhibits a peak near the maximal spectral amplitude. Within the same range of the control parameter, the width of the spectral peak reaches a minimum.

**Figure 3.**
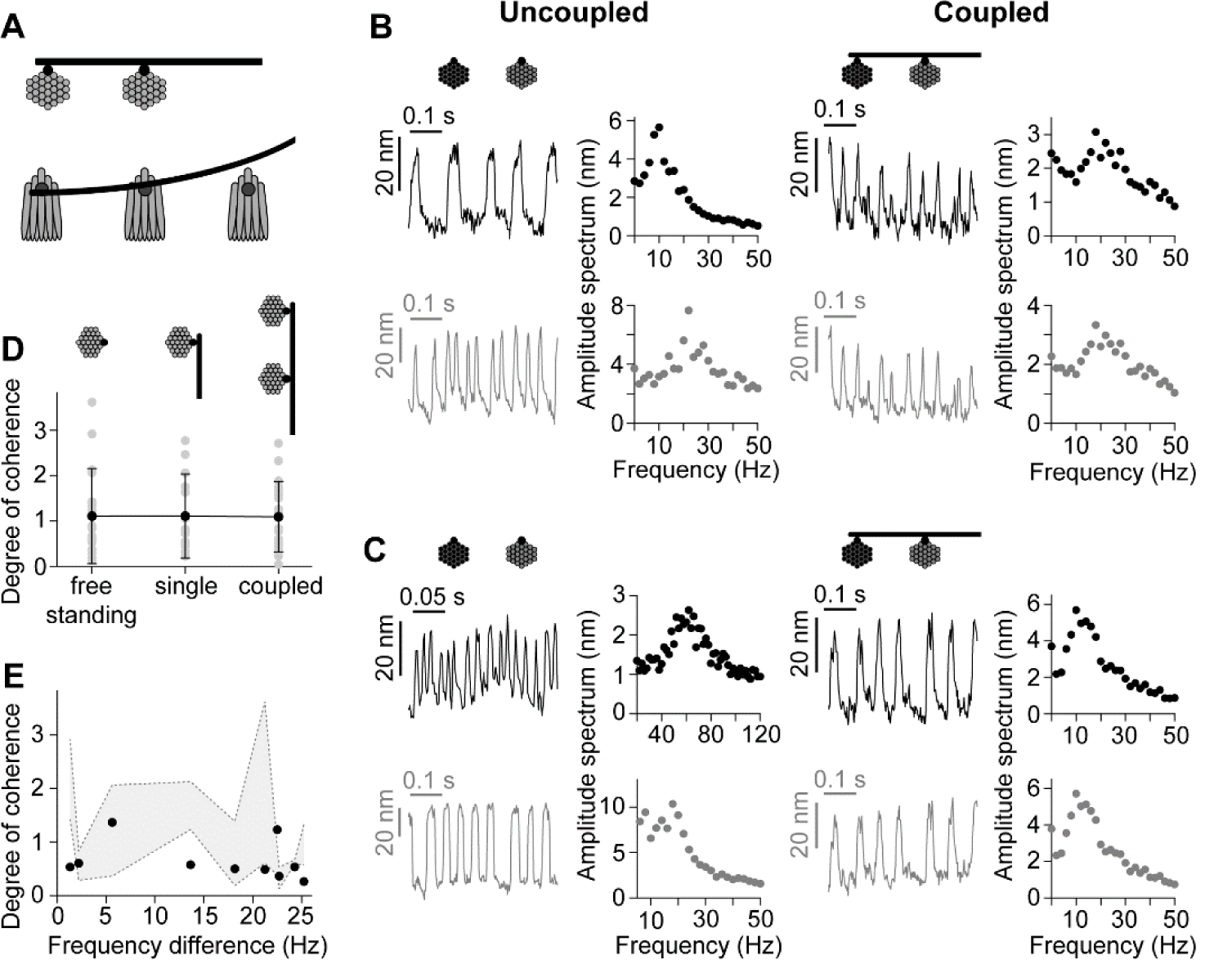
(*Experiments*) Synchronized spontaneous oscillations of hair bundles. A) A schematic drawing depicts the coupling of two hair bundles by connecting a glass fiber, observed from the top-down view of the epithelium. A side view diagram illustrates the curvature of the fiber that allows us to avoid contacting other nearby hair bundles. B)-C) (Left panels, ‘Uncoupled’) Two examples of spontaneous oscillations of two pairs of hair bundles in the absence of coupling. (Right panels, ‘Coupled’) Upon coupling, the displacements of the two bundles become nearly identical. The corresponding amplitude spectra are shown to the right of the time traces. Error bars are the standard deviations from the finite-time Fourier transform. D) The average degree of coherence (black dots) of hair bundles under three types of mechanical manipulations: unloaded (‘free-standing’), loaded without coupling to another bundle (‘single’), and coupled bundles (‘coupled’) are 1.11±0.94 (n=20), 1.11±0.82 (n = 15), and 1.09±0.69 (n= 26), respectively. Error bars indicate standard deviation of the data from individual hair bundles (grey dots). E) The degree of coherence of synchronized oscillations (black dots) is independent of the bundles’ characteristic frequencies or their regularity in the absence of coupling. The degree of coherence of the two uncoupled hair bundles are illustrated by the upper and the lower bounds of the grey area.

For part of this study, we included complex gaussian white noise terms *η*_*j*_(*t*) = *η*_*j,R*_(*t*) + *iη*_*j,I*_(*t*), where *j* = 1,2, which satisfied ⟨*η*_*j,α*_(*t*)*η*_*k,β*_(*t′*)⟩_*t*_ = *σ*^2^*δ*(*t* − *t′*)*δ*_*jk*_*δ*_*αβ*_, where ⟨… ⟩_*t*_ denotes time average, *j, k* =1,2, and *α, β* denotes *R* or *I*. The noise variance, *σ*^2^, was adjusted such that the ratio between the root-mean-square displacement to the spectral amplitude approximately matched the experimental range. Numerical simulations of Eqs. 1 - 2 were performed using the 4^th^-order Runge Kutta method at a 10-ms time step. The initial conditions of both oscillators were chosen to be real and identical at 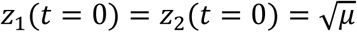.

### Biological preparation and measurement of hair bundle displacement

All animal-handling protocols were approved by the University of California Los Angeles Chancellor’s Animal Research Committee (Protocol Number ARC 2006-043-41A). Sacculi were excised from the inner ears of North American bullfrog (*Lithobates catesbeianus*). The preparation was mounted into a two-compartment chamber in which the basal side was immersed in artificial perilymph (110 mM Na^+^, 2 mM K^+^, 1.5 mM Ca^2+^, 113 mM Cl^-^, 3 mM d-glucose, 1 mM sodium pyruvate, 1 mM creatine, and 5 mM HEPES), and the apical side in artificial endolymph (2 mM Na^+^, 118 mM K^+^, 0.25 mM Ca^2+^, 118 mM Cl^-^, 3 mM d-glucose, and 5 mM HEPES). The otolithic membrane was gently removed after 8-min enzymatic dissociation with 50 μg/mL Collagenase IV (Sigma Aldrich, St. Louis, MO). For experiments in which calcium concentration of endolymph was varied, the preparation was initially immersed in low-calcium endolymph with 0.1 mM Ca^2+^. The calcium level was increased by increments of 75 μM by adding 5-μL of CaCl2 containing 100 mM Ca^2+^.

Images of the preparation were recorded at 500 frames/second by a high-speed CMOS camera (Orca-Flash4.0, Hamamatsu, Shizuoka, Japan). The hair bundle motion was tracked with a custom software written in MATLAB (The MathWorks, Natick, MA). The center position of the bundle was extracted by calculating the center of gravity of the intensity profile in each frame. A slow drift in the bundle displacement was removed by applying a 2^nd^-order Savitzky-Golay filter to each 5-s segment of the recording.

### Mechanical coupling of two hair bundles and stimulation

A borosilicate glass probe was pulled with a micropipette puller (P97, Sutter Instruments, Novato, CA). An additional rod of approximately 1-μm diameter was fabricated at a 90° angle from the tip of the probe using a microforge. The curvature of the rod was then created by placing the probe’s tip near the microforge. The difference in the temperatures of the two sides of the rod resulted in a bending away from the heat source. This structure allowed us to attach the probe’s tip to the kinociliary bulbs of two hair bundles, while avoiding other hair bundles on the preparation (Fig. 3A). To calibrate the stiffness of the glass fiber, the motion of its tip in water was imaged at 10,000 frames/s, and its power spectral density was fitted by a Lorentzian function. The glass fibers utilized in this work had stiffnesses of ∼0.1 – 0.2 mN/m.

To deliver sinusoidal stimulation at various frequencies, the base of the probe was mounted on a piezoelectric stimulator (P-150, Piezosystem Jena, Hopedale, MA) and controlled with LabVIEW (National Instruments, Austin, TX). The probe base was driven by a discrete frequency sweep sinusoidal signal at 30 nm amplitude. This corresponded to a maximal force of ∼3 – 6 pN on a stationary hair bundle. The frequency was increased from 2.5 Hz to 50 Hz at 2.5-Hz increments. Each stimulus frequency was presented for 20 cycles.

### Data analysis

All analyses were performed in MATLAB. Time traces obtained from numerical simulations and hair bundle displacement measurements from experiments were analyzed in a similar fashion. The oscillation amplitudes and frequencies were extracted by two methods. First, the spectral amplitude and spectral frequency were determined from the parameters of the Lorentzian fit to the amplitude spectrum. We calculated the finite-time Fourier transform on each nonoverlapping segment of the signal. The duration of each segment was 1 second for experimental recordings and 50 seconds for simulated traces. The amplitude spectrum was the averaged magnitude of the complex transform. For the second approach, we identified the local extrema of oscillations using a peak detection algorithm. All local extrema were determined from a trace smoothed with a five-point moving average. The threshold for oscillation detection was defined to be 20% of the difference between the global maximum and minimum, to exclude random fluctuations from the analysis. Any local maximum (minimum) whose displacement exceeded the adjacent minimum (maximum) by more than the threshold was identified as a peak (trough). The peak amplitude was computed by averaging half the difference between any adjacent peak and trough for the entire duration of the analyzed signal. Similarly, the peak-to-peak frequency was determined from the averaged reciprocal of the period between two neighboring peaks.

We quantified the degree of coherence of a spontaneous oscillation from the quality factor – the ratio between its peak spectral frequency and the full width at half maximum of the peak in the amplitude spectrum. In response to sinusoidal driving forces at various frequencies, the phase-locked amplitude was estimated from a sinusoidal fit to the average bundle displacement. A Lorentzian function was fitted to the plot of the response amplitude as a function of the driving frequency to extract the quality factor.

To compute the vector strength of the phase difference, the instantaneous phase of each time trace, denoted by *ϕ*_*j*_(*t*), where *j* = 1,2, was obtained from the Hilbert transform. The vector strength was defined as *V* = |⟨*e*^*i*Δ*ϕ*(*t*)^⟩_*t*_|, where Δ*ϕ*(*t*) = *ϕ*_1_(*t*) − *ϕ*_2_(*t*) denotes the oscillators’ phase difference at time *t*. Perfectly phase-locked signals had a vector strength of unity, whereas uncorrelated signals had zero vector strength.

Hartigan’s dip statistic was used to indicate the crossing of a bifurcation of a noisy nonlinear system by quantifying the multimodality of position distributions. The algorithm measured the statistical difference between the empirical distribution and a uniform (null) distribution [29]. A noisy motion underlaid by a limit-cycle oscillation possessed a large value of the dip statistic, whereas random fluctuations around a stable fixed point corresponded to a small dip statistic. The two behaviors were distinguished by the value of the dip statistic with a threshold at 0.01 [13].

A two-dimensional phase space was reconstructed from experimental recordings, to investigate the dynamical trajectory of the oscillator. The time trace was represented as the real part of an analytical signal whose imaginary part was obtained from the Hilbert transform of the oscillation [13]. The distribution of the trajectory in the complex plane, termed the analytic distribution, was created from a two-dimensional histogram of the analytic signal. An analytic distribution with a loop structure corresponded to a limit-cycle oscillation, whereas a unimodal distribution corresponded to random fluctuations around a stable fixed point.

To estimate the critical control parameter *μ*_*c*_ of coupled oscillators from numerical simulations of the model, the initial positions of both oscillators were fixed at an identical, small value ∼10^−4^ – 10^−3^. The oscillation amplitude decayed towards zero when poised in the stable regime, *μ* < *μ*_*c*_, or grew towards a constant value corresponding to the radius of the limit-cycle in the oscillatory regime, *μ* > *μ*_*c*_. At the bifurcation, *μ* = *μ*_*c*_, the duration of the transient solution diverged, and the oscillation profile remained relatively unchanged regardless of the initial condition. We extracted the time-dependent peak amplitude of the transient solution simulated at different values of *μ* and plotted its rate of change as a function of *μ*. The critical value, *μ*_*c*_, was identified at the zero crossing. We increased the control parameter at 0.005-increments, which determined the resolution of the estimated *μ*_*c*_. For the parameter space explored in this work, the critical values *μ*_*c*_ of both oscillators were identical within the 0.005 resolution.

## RESULTS

### Theoretical predictions

A system of two hair bundles connected to a glass fiber was modeled as two mutually coupled nonlinear oscillators (Methods). In the absence of coupling, each oscillator displayed a transition between a fixed point and limit-cycle behavior via a supercritical Hopf bifurcation at a critical control parameter equal to zero. The dynamics of the oscillators were described by the complex variables, *z*_1_(*t*) and *z*_2_(*t*), with their real parts representing the oscillators’ displacements. *ω*_0,1_ and *ω*_0,2_ denote the characteristic frequencies, with *ω*_0,2_ ≥ *ω*_0,1_.

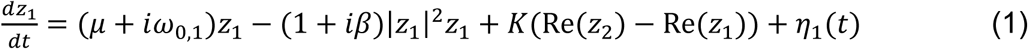

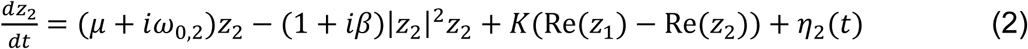

The coupling was modeled as an elastic element whose extension was the relative displacement of the oscillators, i.e., the difference in the real parts of *z*_1_(*t*) and *z*_2_(*t*). To describe the strong coupling conditions experienced by the hair bundles connected by a short glass fiber, the coupling constant was set to be real and denoted by a stiffness *K*. Our model thus incorporated coupling only between the real components of the dynamical variables.

#### Critical control parameter of coupled oscillators

We first identified the critical control parameter *μ*_*c*_ of coupled oscillators under different coupling constants *K. μ*_*c*_ was estimated from the rate of change in the time-dependent peak amplitude of the transient oscillation (Methods). The rate became zero at criticality, *μ* = *μ*_*c*_. Within the range of parameters explored in our study, *μ*_*c*_ of both oscillators remained nearly identical. As the stability of the system near the bifurcation point does not involve the nonlinear term, simulations were performed in the isochronous limit, *β* = 0, for simplicity.

When the oscillators’ characteristic frequencies were nearly identical, *ω*_0,1_ ≈ *ω*_0,2_, *μ*_*c*_ remained close to zero and was unaffected by the coupling constant (Fig. 1A). With the introduction of a frequency difference, the magnitude of *μ*_*c*_ increased over a range of low coupling constants, displayed a maximum, and declined towards zero at higher stiffnesses. The peak *μ*_*c*_ occurred at progressively larger coupling strengths as the characteristic frequencies diverged. The root-mean-square displacement, calculated from the steady-state solution of each oscillator, confirmed the existence of a quiescent regime for control parameters below *μ*_*c*_ (Fig. 1B).

When the control parameters were fixed at a sufficiently large value, the self-sustained motion displayed higher-order mode locking at low coupling coefficients (Fig. 1C). A slight increase in the stiffness of the coupling element suppressed the oscillations, as *μ*_*c*_ exceeded the control parameter of each oscillator. Synchronized limit-cycle oscillations then emerged again at high coupling constants, with unity vector strength of their phase difference (Fig. 1, B and C). These results indicated that, in the presence of a large frequency difference, oscillators individually poised in the oscillatory regime may be brought near a critical point or into the quiescent regime by adjusting the strength of coupling. Further, the small value of *μ*_*c*_ at high stiffnesses suggested that strongly coupled oscillators could display synchronized autonomous oscillations despite a large difference in their characteristic frequencies.

To elucidate the dynamics of coupled oscillators in the complex phase space, we investigated the phase portrait constructed from the real and imaginary part of *z*_1_(*t*) and *z*_2_(*t*). The analysis revealed that the shift in *μ*_*c*_ was associated with an ‘amplitude death’ regime [30, 31]. For *μ* > *μ*_*c*_, an isolated oscillator displayed an unstable fixed point at the origin of the complex phase space. This fixed point became stable in the presence of coupling with an appropriate stiffness coefficient. Both phase space trajectories were attracted to the origin, rendering the suppression of autonomous motion (Fig. 1C). Upon an increase in the coupling constant, the fixed point lost its stability, and limit cycles emerged. Note that the imaginary components of the two trajectories were not necessarily identical, as the coupling only affected the real parts of the dynamical variables.

A linear stability analysis of Eqs. 1 – 2 showed that, for nearly identical oscillators, *ω*_0,2_/*ω*_0,1_ *≃* 1, a supercritical Hopf bifurcation occurred at *μ*_*c*_∼(*ω*_0,2_ − *ω*_0,1_)^2^/4*K* (Supporting material, II). On the other hand, at a sufficiently large frequency difference, *ω*_0,2_/*ω*_0,1_ ≳ 2, *μ*_*c*_ satisfied

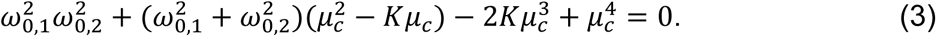

Numerical solutions of *μ*_*c*_ as a function of *K* from Eq. 3 agreed with results from simulations, verifying consistency between the simulations and analytic estimates (Figs. 1, A – B and S1).

To facilitate comparisons to the experimental data, we focused on the strong-coupling limit, in which the stiffness was fixed at a large value (Supporting Material, I); the coupling coefficient was kept constant at *K* = 100 for the rest of our study. In this limit, *μ*_*c*_ displayed a monotonic increase with the frequency difference, but always remained closed to zero (Fig. S1). The oscillation amplitude obtained from numerical simulations of the model was proportional to (*μ* − *μ*_*c*_)^1/2^, characteristic of a system near a supercritical Hopf bifurcation (Fig. 1D). We note that, in this strong coupling limit, nearly identical oscillators with appropriate initial positions could display an oscillation death behavior when *μ* > *μ*_*c*_ (Supporting Material, III).

#### Dynamics of synchronized oscillations far from a bifurcation

Next, we investigated the behavior of synchronized limit-cycle oscillations far from criticality, *μ* > *μ*_*c*_. We focused on this regime because free-standing hair bundles from the bullfrog sacculus typically display robust large-amplitude spontaneous oscillations under *in vitro* conditions, and thus seem to be innately poised far from the quiescent regime.

In the oscillatory regime, far from the bifurcation, the oscillators displayed two classes of behavior, dependent on their degrees of nonisochronicity *β*. For small *β*, synchronized oscillations displayed similar features to those of isochronous (Fig. 1D) or uncoupled oscillators (Fig. S4). The peak amplitude of the oscillation rose monotonically as the control parameter was increased (Fig. 2A). Due to the positive *β*, the peak-to-peak oscillation frequency declined upon growth of the amplitude.

When *β* was sufficiently large, the limit-cycle amplitude displayed a broad peak with the maximum occurring between control parameter values 0.5 and 1.0. For oscillators with a small frequency difference, the tuning was due to the suppression of oscillations within a narrow range of control parameter values between 0.9 and 1.1 (Fig. 2D). On the other hand, oscillators with a large frequency difference exhibited the maximal amplitude that approximately coincided with the minimum in the oscillation frequency (Fig. 2G).

The phase portraits constructed from the real and imaginary parts of *z*_1_(*t*) and *z*_2_(*t*) revealed that the tuning behavior of highly nonisochronous oscillators was strongly influenced by their angular nullclines in the uncoupled limit, which coincided with the limit cycle radii when the control parameter was *μ*_0,1_ or *μ*_0,2_. In polar coordinates, the angular equation for the *j*^th^ oscillator, in the absence of coupling, was given by 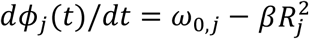, where 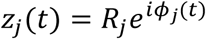. The angular nullcline resided where the oscillation frequency vanished. This coincided with a limit cycle whose radius satisfied 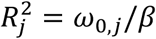 when the control parameter was *μ*_0,*j*_ = *ω*_0,*j*_/*β*.

In contrast to nearly identical oscillators, which exhibited similar trajectories in the phase portraits (Figs. 2D and S2), oscillators with a large frequency difference displayed stable limit cycles with drastically distinct shapes (Fig. 2A and 2G). The first oscillator’s trajectory was approximately elliptical and situated on the upper perimeter of the second oscillator’s limit cycle. This behavior exhibited a bistability in which the first trajectory could instead be confined to the lower half of the complex plane.

For small *β*, the range of control parameters investigated did not approach the angular nullclines, thus resulting in the monotonically varying limit cycle amplitudes and frequencies of both oscillators (Fig. 2D). For large *β* and small frequency difference, the limit cycle trajectories terminated at two distinct points in the vicinity of the angular nullclines (Fig. 2D), indicative of an oscillation death regime (Supporting material, III). This quiescent behavior gave rise to a local maximum in the limit-cycle amplitude within a range *μ*_*c*_ < *μ* < *μ*_0,1_ ≈ *μ*_0,2_. The system displayed stable, synchronized limit cycles at higher values of the control parameter. In contrast, oscillators with a large frequency difference reached the peak in their oscillation amplitude and a minimal frequency when the trajectory of the first oscillator approached its angular nullcline. A larger control parameter led to the separation of the two limit cycles as well as an attenuation of the oscillation amplitude (Fig. 2G).

Finally, we introduced Gaussian white noise into Eqs. 1 – 2 to account for the noisy nature of hair bundles’ motion. The features of noisy oscillations were extracted from the amplitude spectra obtained from a finite-time Fourier transform. As in the deterministic case, the dynamics of the oscillators were consistent with a supercritical Hopf bifurcation in the transition regime (Fig. S3) and possessed two classes of behaviors far from the critical point (Fig. 2, B, E and H).

We quantified the coherence of the oscillations by the ratio of the spectral frequency to the full width at half the maximum of the peak in the amplitude spectrum (Methods). Near criticality, the degree of coherence was enhanced as the oscillators crossed the bifurcation point and entered the oscillatory regime. As the control parameter continued to increase, the degree of coherence reached a plateau for weakly nonisochronous oscillators (Fig. 2C), whereas a nonmonotonic behavior was observed in oscillators with a large *β*. For nearly identical oscillators, the vanishing oscillation frequency or amplitude led to a minimal degree of coherence over a broad range of the control parameter (Fig. 2F). Hence, a peak coherence was observed close to the critical point. In contrast, the motion of oscillators with a large frequency difference became more regular as the oscillation amplitude increased. Therefore, the degree of coherence, as well as the amplitude, reached their maxima over the same range of control parameters (Fig. 2I).

### Experimental observations

We attached the tip of a thin, bent glass fiber to two neighboring oscillatory bundles in the planar epithelium of the bullfrog sacculus and observed their dynamics under high coupling stiffness (Supporting material, I). The curvature of the tip of the fiber enabled us to avoid other hair bundles on the preparation (Fig. 3A).

#### Synchronized spontaneous oscillations of coupled hair bundles

When immersed in an artificial endolymph with 250-μM calcium, the concentration near the physiological level, coupled hair bundles oscillated spontaneously with a pronounced peak in the amplitude spectrum (Fig. 3 B and C). The bundles always moved in synchrony with the vector strength of their phase difference exceeding 0.9. These synchronized spontaneous oscillations were observed despite the large difference in their natural frequencies measured in the absence of an attached fiber.

We studied the effects of coupling on the fluctuation levels by extracting the degree of coherence of coupled and uncoupled hair bundles. As the effects of thermal noise depended greatly on the geometry and stiffness of the system, we compared the regularity of coupled hair bundles’ oscillations to those of single bundles, each attached to a fiber with comparable stiffness and drag coefficients. As hair bundles could sustain microscopic damage upon removal of the fiber, thus affecting their dynamics, these comparisons were performed on data sets obtained from two distinct groups of hair cells. We found that the average degrees of coherence in both cases were comparable (Fig. 3D): 1.11±0.82 (n = 15) for single bundles and 1.09±0.69 (n= 26) for coupled bundles. In addition, the degree of coherence was not significantly different from free-standing bundles with no fiber attached: 1.11±0.94 (n=20).

We recorded the motion of pairs of free-standing hair bundles prior to the coupling and extracted their degree of coherence. We found that, in contrast to previous predictions [21, 25], neither a small frequency difference, nor a high regularity of the uncoupled oscillations improved the coherence of the synchronized motion (Fig. 3E). This contradiction could partly stem from the variations in the degree of nonisochronicity of the bundles. Our theoretical predictions suggested that the coherence of synchronized oscillations that were far from the bifurcation depended on the frequency difference and the coefficient *β* in a complex manner (Fig. 2, C, F, and I).

#### Coupled hair bundle motility near the critical point

Next, we employed the variation of calcium concentration in the endolymph as an experimental modulation of the control parameter. The level of calcium was increased from 100 μM in 75-μM increments (Methods). In this section, we focus on the dynamics of hair bundles at high calcium concentrations, near the critical level (Fig. 4, shaded areas). The behavior of spontaneous oscillations at lower calcium levels will be discussed in the next section.

**Figure 4.**
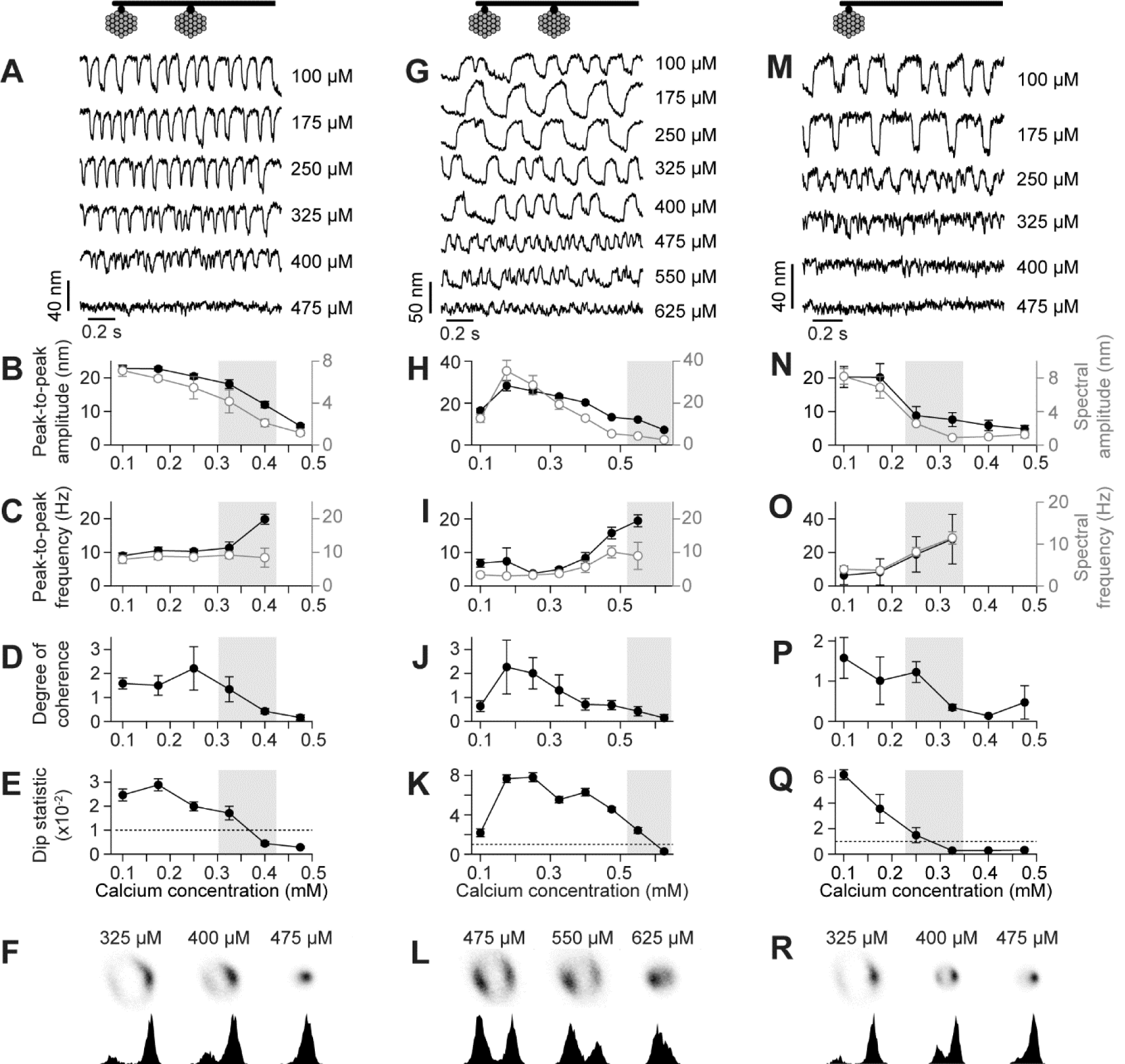
(*Experiments*) Synchronized spontaneous oscillations under calcium manipulation. A) Coupled hair bundles’ oscillations at different calcium levels, indicated to the right of each trace. In B)-E), shaded areas indicate the transition regime from oscillatory to quiescent behavior. B) The peak (black dots) and spectral amplitude (gray open dots) monotonically decrease as calcium level is elevated. C) The peak-to-peak (black dots) and spectral frequency (gray open dots) remain relatively unchanged. In the transition regime, the peak-to-peak frequency increases, whereas the spectral frequency is poorly determined. At a higher calcium level, the hair bundle motion becomes indistinguishable from a random fluctuation, and the oscillation frequency cannot be extracted. D) The degree of coherence of spontaneous oscillations is minimally affected at lower calcium levels and gradually declines in the transition regime. E) The Hartigan’s dip statistic, in association with the oscillation amplitude, decreases monotonically with increasing calcium levels. F) The analytic distribution (top panels), as well as the displacement histogram (bottom panels), show a transition from a ring-like structure to a unimodal distribution. In the transition regime, the oscillation spends longer fractions of time deflected in the positive direction. G) Another type of behavior is observed from coupled hair bundles whose oscillations become larger at 175-250 μM calcium. The calcium levels are indicated to the right of each trace. H)-L) A peak in the oscillation amplitude, degree of coherence, and the dip statistic are observed near the physiological calcium level. M) An example of a single bundle’s displacement loaded by a glass fiber, at various calcium levels indicated to the right of each trace. N)-R) The oscillation behaviors qualitatively resemble the first type of coupled hair bundles (A)-F)). In all cases, coupled or uncoupled hair bundles display similar behaviors in the transition regime (shaded areas). Error bars indicate the standard deviation of the data extracted from 6 time trace segments; each is 10 seconds.

We utilized two metrics previously developed to study the behavior of noisy hair bundle motility near the transition between the oscillatory and quiescent regimes (Methods) [13]. Hartigan’s dip statistic was used to identify the crossing of a bifurcation by measuring the degree of multimodality in the distribution of a bundle’s position. We further characterize the dynamics of hair bundle oscillations by reconstructing an analytical signal whose two-dimensional histogram could reveal an underlying stable limit cycle or a stable fixed point in the complex phase space.

The critical range of calcium concentration was indicated by the Hartigan’s dip statistic crossing the threshold of multimodality (Fig. 4, E and K). This signified crossing of a bifurcation: an increase in the calcium level corresponded to a decrease in the control parameter from an oscillatory regime towards a quiescent regime. As the concentration approached a critical range, typically exceeding ∼300 μM, hair bundle motion could become highly asymmetric with the oscillations spending more time in the positive or negative displacement (Fig. 4, A and G). Due to the non-sinusoidal oscillation profile, we extracted the oscillation amplitude and frequency from both the amplitude spectrum and local extrema in the time trace. As the calcium level increased, the spectral and peak amplitudes of synchronized spontaneous oscillations were attenuated, until they became indistinguishable from random fluctuations (Fig. 4, B and H). In association with this, the peak-to-peak frequency increased and became undefined as the oscillations were suppressed (Fig. 4, C and I). The spectral frequency, on the other hand, remained relatively constant as the spectral peak was predominantly broadened, rendering the determination of the peak frequency less reliable.

The analytic distribution in a two-dimensional phase space formed a ring-like structure at low calcium concentrations. The ring decreased in size and converged to a unimodal distribution upon increasing the calcium concentration (Fig. 4, F and L). We note that at the levels of calcium concentration employed in this study, we did not observe an analytical distribution that corresponded to a coexistence of a limit-cycle and a stable fixed point at the origin. Our results predominantly indicated that the behavior of coupled hair bundles across the critical calcium concentration was consistent with a supercritical Hopf bifurcation. Qualitatively identical dynamics were observed in single hair bundles under similar mechanical loading (Fig. 4, M – R).

#### Tuning behavior of synchronized spontaneous oscillations

In this section, we investigated the dynamics of hair bundles at lower calcium levels, far from criticality, and compared them to the theoretical predictions of coupled oscillators far from a supercritical Hopf bifurcation (Fig. 2).

We observed two types of behaviors of synchronized spontaneous oscillations at low calcium levels. First, some coupled hair bundles showed a monotonic decrease in their oscillation amplitude and the degree of coherence as the calcium was elevated, predominantly consistent with a system of coupled, weakly non-isochronous oscillators (Fig. 4, B – D). The oscillation frequency, however, could remain largely unaffected (Fig. 4C) or continually increase (Fig. S5). On the other hand, the motion of some hair bundles displayed maximal amplitude and coherence at an optimal calcium concentration, typically near the physiological level of 250 μM (Figs. 4, H – J and S5). Within the same range of calcium concentrations, the peak-to-peak frequency of the oscillation clearly exhibited a minimum. This type of behavior agreed with a system of oscillators with a high degree of nonisochronicity.

To examine the frequency selectivity of coupled hair bundles at different calcium levels, we measured the bundles’ displacements in response to pure tone stimulation. Sinusoidal signals at various frequencies were delivered to the base of the glass fiber at a constant amplitude (Methods). The quality factor of the phase-locked amplitude displayed a similar calcium dependence as that shown by the degree of coherence of spontaneous oscillations (Fig. S6).

## DISCUSSION

We observed synchronization between the autonomous motion of two strongly coupled hair bundles from the bullfrog sacculus. Our experimental findings were well reproduced by a theoretical model of coupled nonlinear oscillators, each displaying a supercritical Hopf bifurcation. Results from the model indicated that strong coupling minimally affected the critical control parameter of the oscillators. This indicated that the oscillatory hair bundles, despite their vastly different natural frequencies, remained in the unstable regime and far from criticality upon coupling, yielding the observed synchronized spontaneous oscillations. The theoretical prediction also implied that the previously proposed mathematical descriptions of uncoupled hair bundles should be applicable to a small number of bundles under strong coupling.

Under weaker coupling, on the other hand, the model demonstrates that oscillators with different characteristic frequencies may be brought closer to a supercritical Hopf bifurcation by a shift in the critical point. Hence, while the control parameter of each individual oscillator is unaffected, the coupling shifts the bifurcation point closer to them. Under physiological conditions, groups of hair bundles in the inner ears of many species experience weak coupling at large spatial separations. Our theoretical prediction thus proposes an additional potential mechanism of tuning a system of oscillatory hair bundles closer to criticality via mechanical coupling. Other proposed physical quantities have focused on poising an uncoupled hair bundle near the verge of an instability; examples include mass, elastic loadings, and a stationary offset on the bundle’s resting position imposed by its surrounding structure [12, 15, 32]. A combination of these may be utilized as a mechanism that effectively brings an *in vivo* system of hair bundles near the critical point.

We utilized the level of calcium concentration in the endolymph as an experimental means of modulating the control parameter across a supercritical Hopf bifurcation. Far from the critical point, theoretical results suggest that individual hair bundles may be regarded as nonisochronous oscillators: the oscillation frequency increases upon an attenuation in the amplitude. An appropriate degree of nonisochronicity, determined by the hair bundles’ natural frequencies, was required for the enhanced regularity of the synchronized spontaneous motion near the physiological calcium level. The nonisochronicity was further shown to enhance the phase-locked amplitude in response to sinusoidal stimuli.

Several studies have demonstrated the nonisochronous behavior of individual hair bundles under different types of manipulations, including a change in calcium concentration, loading by a mass or an elastic element, and an application of a constant force [15, 33]. A previous investigation showed that a high degree of nonisochronicity may be beneficial to the detection of rapidly varying signals by poising the system in a chaotic regime in the presence of noise, and thus enhancing the temporal resolution [34]. Further, a lowered detection threshold was observed at an optimal level of nonisochronicity. Our results indicate that nonisochronicity also affects the dynamics of coupled bundles.

### Possible mechanism underlying hair bundle nonisochronicity

Spontaneous oscillations of hair bundles arise from the gating of the mechanosensitive ion channels, whose open probability is regulated by the tension in a tip link, a thin filament which serves as a component of the gating spring. Several microscopic models of hair bundle’s mechanics have shown that the oscillation amplitude and frequency crucially depend on the stiffness of the gating spring, which dictates the nonlinearity of the system [11, 12, 35]. Further, alterations in the spontaneous oscillation profiles under various forms of mechanical or chemical manipulation are well reproduced by models in which the gating spring stiffness is reduced upon an increase in external calcium [33].

To test whether the calcium-sensitive gating spring stiffness may contribute to hair bundles’ nonisochronicity, we performed numerical simulations of the model previously shown to describe individual hair bundle motility [33] and observed the spontaneous oscillations at different levels of calcium sensitivity. The model postulated that the gating spring incorporated a calcium-binding site whose binding probability *p*_*gs*_ depended on the local calcium concentration [Ca^2+^]_*gs*_, and followed the standard rate equation *dp*_*gs*_/*dt* = *k*_*gs,on*_[Ca^2+^]_*gs*_(1 − *p*_*gs*_) − *k*_*gs,off*_*p*_*gs*_. The rates of binding and unbinding of calcium to the gating spring are denoted by *k*_*gs,on*_ and *k*_*gs,off*_, respectively. The gating spring stiffness *K*_*gs*_ was governed by *K*_*gs*_ = *K*_*gs*,0_ − *K*_*gs*,1_*p*_*gs*_, in which *K*_*gs*,0_ and *K*_*gs*,1_ are positive constants.

We found that the spontaneous oscillations exhibited by a hair bundle with a constant gating spring stiffness, *K*_*gs*,1_ = 0, were minimally dependent on the calcium concentration (Fig. 5A). The autonomous motion abruptly diminished at a critical calcium level. When the gating spring stiffness was modulated by calcium, the oscillation amplitude was attenuated, and the frequency increased upon a rise in the calcium level, a feature that is consistent with a nonisochronous nonlinear oscillator. At a higher calcium sensitivity (*K*_*gs*,1_/*K*_*gs*,0_), the hair bundle displayed a stronger nonisochronous behavior: its oscillation frequency more steeply depended on the amplitude as the calcium concentration increased (Fig. 5, B – D). These results suggest that a calcium-sensitive elastic element may partly account for the hair bundle’s nonisochronicity.

**Figure 5.**
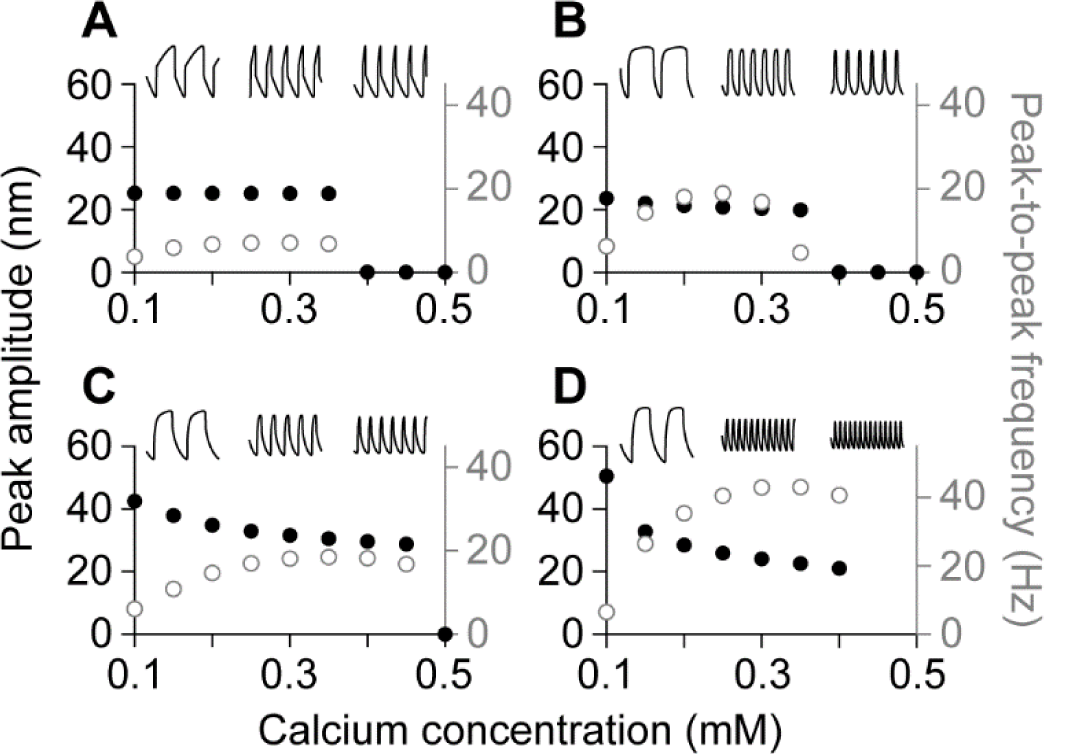
(*Model*) Nonisochronicity of a single hair bundle with a calcium-sensitive gating spring stiffness. Results are obtained from numerical simulations of a model of hair bundle motility. The parameter values are shown in Table S1. In all panels, the peak amplitude (black dots) and the peak-to-peak frequency (gray open dots) of hair bundle spontaneous oscillations at different calcium concentrations are shown. The insets illustrate the time traces of spontaneous oscillations at calcium concentrations 0.1, 0.2, and 0.3 mM (from left to right). A) When the gating spring stiffness is a constant, with *K*_*gs*,0_ = 1050 μN/m, and *K*_*gs*,1_ = 0, the oscillation profiles are not sensitive to calcium. B) For a small calcium sensitivity, with *K*_*gs*,0_ = 1300 μN/m, and *K*_*gs*,1_ = 500 μN/m (*K*_*gs*,1_/*K*_*gs*,0_ = 0.38), the oscillation frequency increases with calcium level, whereas the amplitude is slightly attenuated. C) The decline in the oscillation amplitude becomes more pronounced at a higher calcium sensitivity of the gating spring, with *K*_*gs*,0_ = 2000 μN/m, and *K*_*gs*,1_ = 1200 μN/m (*K*_*gs*,1_/*K*_*gs*,0_ = 0.6). D) At *K*_*gs*,0_ = 2550 μN/m, and *K*_*gs*,1_ = 2000 μN/m (*K*_*gs*,1_/*K*_*gs*,0_ = 0.78), the nonisochronous behavior is strong, with the oscillation frequency rapidly increases upon an attenuation in amplitude.

## CONCLUSION

We investigate a model of two nonisochronous oscillators, whose real parts of the complex dynamical variables are mutually coupled. Results from the model suggest an alternative mechanism that tunes a system of weakly coupled unstable oscillators closer to criticality via a shift of the bifurcation point. Theoretical predictions for strongly coupled oscillators are consistent with the dynamics of hair bundles under strong mechanical coupling. Within a critical range of calcium concentration in the endolymph, synchronized spontaneous oscillations of hair bundles become suppressed and exhibit characteristics of a nonlinear system crossing a supercritical Hopf bifurcation. At calcium levels sufficiently far from the critical point, hair bundles may display a tuning in the amplitude and coherence of their spontaneous oscillations, as well as a tuning of the quality factor in response to an external driving force. This behavior was observed in the model with highly nonisochronous oscillators. Further, results from a microscopic model of a single hair bundle motility suggest that nonisochronicity may stem from a calcium-sensitive elastic element that controls the gating of the mechanosensitive ion channels.

## AUTHOR CONTRIBUTIONS

Y.R. designed the research, conducted physiological recordings and theoretical analysis, and wrote the manuscript. J.F. designed the research, performed theoretical analysis, and wrote the manuscript. D.B. supervised the research, and wrote the manuscript.

## ACKNOWLEDGMENTS

D.B. gratefully acknowledges support of NSF Biomechanics and Mechanobiology Program, under grant 1916136. Y.R. is supported by Chulalongkorn University: CU_GR_62_71_23_28.

